# Chromothripsis-like patterns are recurring but heterogeneously distributed features in a survey of 22,347 cancer genome screens

**DOI:** 10.1101/001776

**Authors:** Haoyang Cai, Nitin Kumar, Homayoun C. Bagheri, Christian von Mering, Mark D. Robinson, Michael Baudis

## Abstract

**Background:** Chromothripsis is a recently discovered phenomenon of genomic rearrangement, possibly arising during a single genome-shattering event. This could provide an alternative paradigm in cancer development, replacing the gradual accumulation of genomic changes with a “one-off” catastrophic event. However, the term has been used with varying operational definitions, with the minimal consensus being a large number of locally clustered copy number aberrations. The mechanisms underlying these chromothripsis-like patterns (CTLP) and their specific impact on tumorigenesis are still poorly understood.

**Results:** Here, we identified CTLP in 918 cancer samples, from a dataset of more than 22,000 oncogenomic arrays covering 132 cancer types. Fragmentation hotspots were found to be located on chromosome 8, 11, 12 and 17. Among the various cancer types, soft-tissue tumors exhibited particularly high CTLP frequencies. Genomic context analysis revealed that CTLP rearrangements frequently occurred in genomes that additionally harbored multiple copy number aberrations (CNAs). An investigation into the affected chromosomal regions showed a large proportion of arm-level pulverization and telomere related events, which would be compatible to a number of underlying mechanisms. We also report evidence that these genomic events may be correlated with patient age, stage and survival rate.

**Conclusions:** Through a large-scale analysis of oncogenomic array data sets, this study characterized features associated with genomic aberrations patterns, compatible to the spectrum of “chromothripsis”-definitions as previously used. While quantifying clustered genomic copy number aberrations in cancer samples, our data indicates an underlying biological heterogeneity behind these chromothripsis-like patterns, beyond a well defined “chromthripsis” phenomenon.

## Background

One consistent hallmark of human cancer genomes are somatically acquired genomic rearrangements, which may result in complex patterns of regional copy number changes [1,2]. These alterations have the potential to interrupt or activate multiple genes, and consequently have been implicated in cancer development [3]. Analysis of genomic rearrangements is essential for understanding the biological mechanisms of oncogenesis and to determine rational points of pharmacological interference [4,5]. Some large-scale efforts have been undertaken to correlate genomic rearrangements to genome architecture as well as to the progression dynamics of cancer genomes [6,7]. At the moment, the stepwise development of cancer with the gradual accumulation of multiple genetic alterations is the most widely accepted model [8].

Recently, using state-of-the-art genome analysis techniques, a phenomenon termed “chromothripsis” was characterized in cancer genomes, defined by the occurrence of tens to hundreds of clustered genomic rearrangements, having arisen in a single catastrophic event [9]. In this model, contiguous chromosomal regions are fragmented into many pieces, via presently unknown mechanisms. These segments are then randomly fused together by the cell’s DNA repair machinery. It has been proposed that this “shattering” and aberrant repair of a multitude of DNA fragments could provide an alternative oncogenetic route [9], in contrast to the step-by-step paradigm of cancer development [8–10]. The initial study reported 24 chromothripsis cases, with some evidence of a high prevalence in bone tumors [9].

Besides human cancers, recent studies have also reported chromothripsis events in germline and non-human genomes [11–13]. However, due to the overall low incidence of this phenomenon, most studies were limited to relatively small numbers of observed events. For example, in a study screening 746 multiple myelomas by SNP arrays, only 10 cases with chromothripsis-like genome patterns were detected [14]. Larger sample numbers are required to gain further insights into features and mechanisms of these events in different cancers.

In contrast to a strict definition of chromothripsis events relying on sequencing based detection of specific genomic rearrangements [15], other studies [7,14,16] have described chromothripsis events based on genomic array analysis without support from whole genome sequencing data. Overall, the minimal consensus of array based studies is the detection of a large number of locally clustered CNA events. In table 1, we provide an overview of studies which so far have reported instances of “chromothripsis” in human cancers [7,9,11,13–14, 16–35].

**Table 1.**
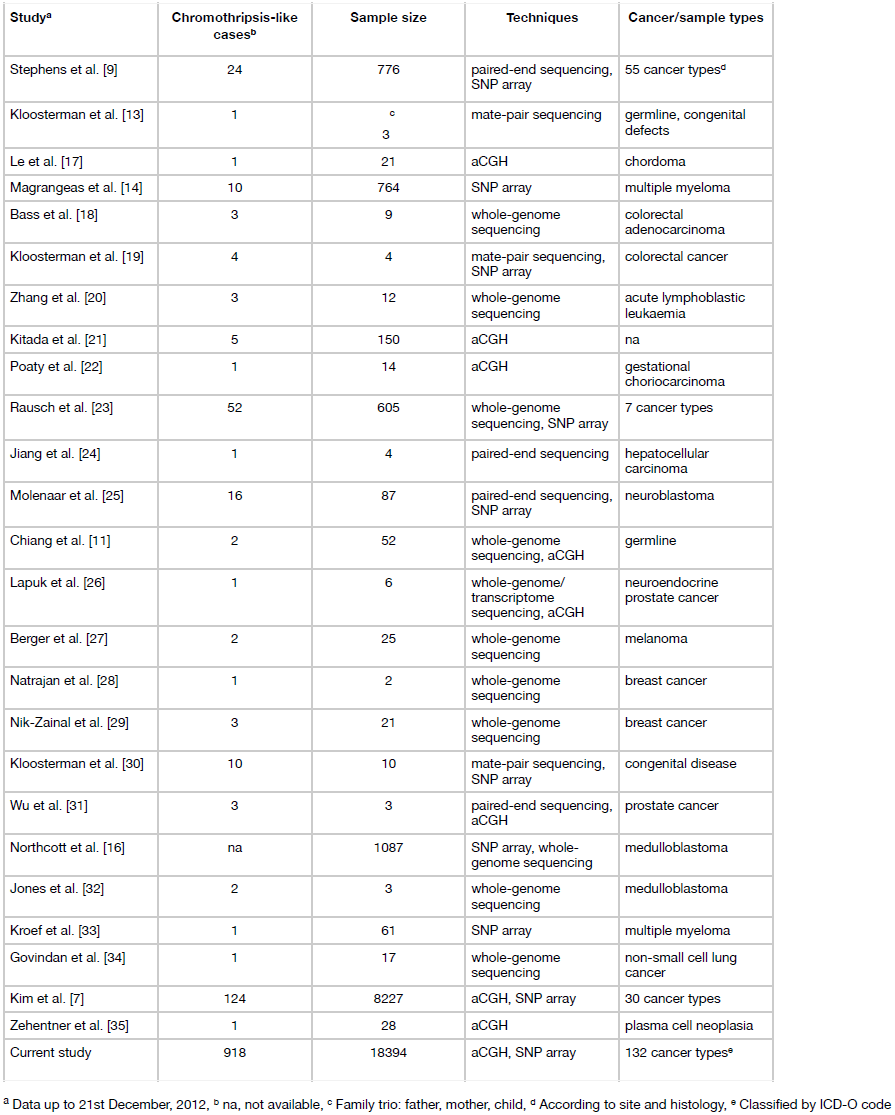
Summary of chromothripsis-like cases identified in previous and current studies.

| Study<sup>a</sup> | Chromothripsis-like cases<sup>b</sup> | Sample size | Techniques | Cancer/sample types |
| --- | --- | --- | --- | --- |
| Stephens et al. [9] | 24 | 776 | paired-end sequencing, SNP array | 55 cancer types<sup>d</sup> |
| Kloosterman et al. [13] | 1 | <sup>c</sup><br>3 | mate-pair sequencing | germline, congenital defects |
| Le et al. [17] | 1 | 21 | aCGH | chordoma |
| Magrangeas et al. [14] | 10 | 764 | SNP array | multiple myeloma |
| Bass et al. [18] | 3 | 9 | whole-genome sequencing | colorectal adenocarcinoma |
| Kloosterman et al. [19] | 4 | 4 | mate-pair sequencing, SNP array | colorectal cancer |
| Zhang et al. [20] | 3 | 12 | whole-genome sequencing | acute lymphoblastic leukaemia |
| Kitada et al. [21] | 5 | 150 | aCGH | na |
| Poaty et al. [22] | 1 | 14 | aCGH | gestational choriocarcinoma |
| Rausch et al. [23] | 52 | 605 | whole-genome sequencing, SNP array | 7 cancer types |
| Jiang et al. [24] | 1 | 4 | paired-end sequencing | hepatocellular carcinoma |
| Molenaar et al. [25] | 16 | 87 | paired-end sequencing, SNP array | neuroblastoma |
| Chiang et al. [11] | 2 | 52 | whole-genome sequencing, aCGH | germline |
| Lapuk et al. [26] | 1 | 6 | whole-genome/transcriptome sequencing, aCGH | neuroendocrine prostate cancer |
| Berger et al. [27] | 2 | 25 | whole-genome sequencing | melanoma |
| Natrajan et al. [28] | 1 | 2 | whole-genome sequencing | breast cancer |
| Nik-Zainal et al. [29] | 3 | 21 | whole-genome sequencing | breast cancer |
| Kloosterman et al. [30] | 10 | 10 | mate-pair sequencing, SNP array | congenital disease |
| Wu et al. [31] | 3 | 3 | paired-end sequencing, aCGH | prostate cancer |
| Northcott et al. [16] | na | 1087 | SNP array, whole-genome sequencing | medulloblastoma |
| Jones et al. [32] | 2 | 3 | whole-genome sequencing | medulloblastoma |
| Kroef et al. [33] | 1 | 61 | SNP array | multiple myeloma |
| Govindan et al. [34] | 1 | 17 | whole-genome sequencing | non-small cell lung cancer |
| Kim et al. [7] | 124 | 8227 | aCGH, SNP array | 30 cancer types |
| Zehentner et al. [35] | 1 | 28 | aCGH | plasma cell neoplasia |
| Current study | 918 | 18394 | aCGH, SNP array | 132 cancer types<sup>e</sup> |

<sup>a</sup> Data up to 21st December, 2012, <sup>b</sup> na, not available, <sup>c</sup> Family trio: father, mother, child, <sup>d</sup> According to site and histology, <sup>e</sup> Classified by ICD-O code

Here, we present a statistical model for the discovery of clustered genomic aberration patterns, similar to those previously labeled as “chromothripsis” events, from genomic array data sets. For the scope of this article, we introduce the term “chromothripsis-like patterns” (CTLP) when discussing those events.

Applying our methodology to 22,347 genomic arrays from 402 GEO (Gene Expression Omnibus) derived experimental series [36], we were able to detect 918 chromothripsis-like cases, and to determine the frequency and genomic distribution of CTLP events in this dataset. Our collection of oncogenomic array data represents 132 cancer types as defined using the ICD-O 3 (International Classification of Diseases for Oncology) coding scheme, enabling us to estimate the incidence of CTLP in diverse tumor types. Among the CTLP cases, varying distributions of fragmented chromosomal regions as well as an abundance of large non-CTLP copy number aberrations (CNA) regions were found, and the genomic context of chromothripsis-like events was investigated. Finally, we evaluated clinical associations of CTLP cottoning samples, based on the clinical information at hand. Overall, this study characterized heterogeneous features of chromothripsis-like events through a large-scale analysis of oncogenomic array data sets and provides a better understanding of clustered genomic copy number patterns in cancer development.

## Results

### Detection of chromothripsis-like patterns from oncogenomic arrays

We collected 402 GEO series, encompassing 22,347 high quality genomic arrays of human cancer samples. A procedure was employed to detect CTLP from these arrays (Figure 1A). The annotated information of the arrays, including normalized probe intensity, segmentation data and quality evaluation, was obtained from our arrayMap database [37] (see Methods for array processing pipeline). After removing technical repeats (e.g. multiple platforms for one sample), a total of 18,394 cases representing 132 cancer types remained. The input data is summarized, at array and case-level, respectively, in Additional file 1: Table S1 and Additional file 2: Table S2. The segmentation data and array profiling can be accessed and visualized through the arrayMap website (www.arraymap.org).

**Figure 1.**
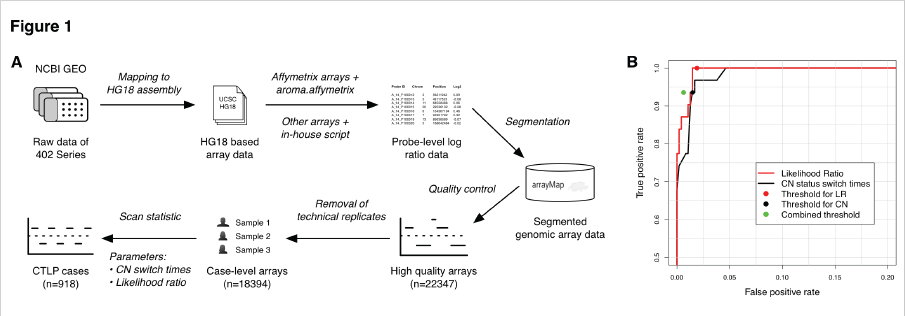
Detection of chromothripsis-like patterns from genomic arrays. (A) Schematic description of the detection procedure. Raw array data of 402 GEO series are first collected and re-analyzed, then annotated and stored in arrayMap database. For high quality arrays, a scan-statistic based algorithm was employed to identify CTLP cases. (B) The ROC curve of the training set and selected thresholds. Two predictors were tested, copy number status change times and the likelihood ratio. Both predictors were integrated into the combined threshold.

According to previous studies, segmental copy number status changes and significant breakpoint clustering are two relevant features of chromothripsis [9,23]. For an automatic identification of CTLP, we developed a scan-statistic based algorithm [38]. We employed a maximum likelihood ratio score, which is commonly used to detect clusters of events in time and/or space and to determine their statistical significance [39] (see Methods). For each chromosome, the algorithm uses a series of sliding windows to identify the genomic region with the highest likelihood ratio as the CTLP candidate. In order to test the performance of the algorithm, 23 previously published chromothripsis cases with available raw array data were collected and used as a training set. This data contained 31 chromothriptic and 475 non-chromothriptic chromosomes that acted as positive and negative controls, respectively (Additional file 3: Table S3). Comparison of copy number status change times and likelihood ratios showed that chromothriptic chromosomes could reliably be distinguished from non-chromothriptic ones (Additional file 1: Figure S1). We generated a receiver operating characteristic (ROC) curve from the training set results, and selected cutoff values based on this curve (copy number status switch times ≥ 20 and log10 of likelihood ratio ≥ 8) (Figure 1B). Furthermore, the sliding window scan statistic accurately identified the genomic regions involved (Additional file 1: Figure S2). Applying this algorithm to the complete input data set, a total of 1,269 chromosomes from 918 cases passed our thresholds and were marked as CTLP events (Additional file 1: Figure S3, Additional file 4: Table S4).

### Chromothripsis-like patterns across diverse tumor types

When evaluating the 1,269 CTLP events, we found a pronounced preference for some chromosomes; this preference showed only limited association with chromosome size (Figure 2A). CTLP occurred more frequently in chromosome 17 than in any other chromosome. This observation is in accordance with data reporting an association between chromothripsis and *TP53* mutations in Sonic-Hedgehog medulloblastoma and acute myeloid leukemia [23]. *TP53* is located in the p arm of chromosome 17, and is involved in cell cycle control, genome maintenance and apoptosis [40,41]. Our dataset showed *TP53* losses in 438 out of 918 (∼48%) CTLP cases, compared to 3,274 out of 17,476 (∼19%) cases in the non-CTLP group (*p* < 2.2×10^−16^; two-tailed Fisher’s exact test; Additional file 2: Table S2). 45 of the 438 *TP53* deletions were part of a CTLP, confirming *TP53* mutation as a recurring event with possible involvement in CTLP formation. Other chromosomes with relatively high incidences of CTLP included chromosomes 8, 11 and 12.

**Figure 2.**
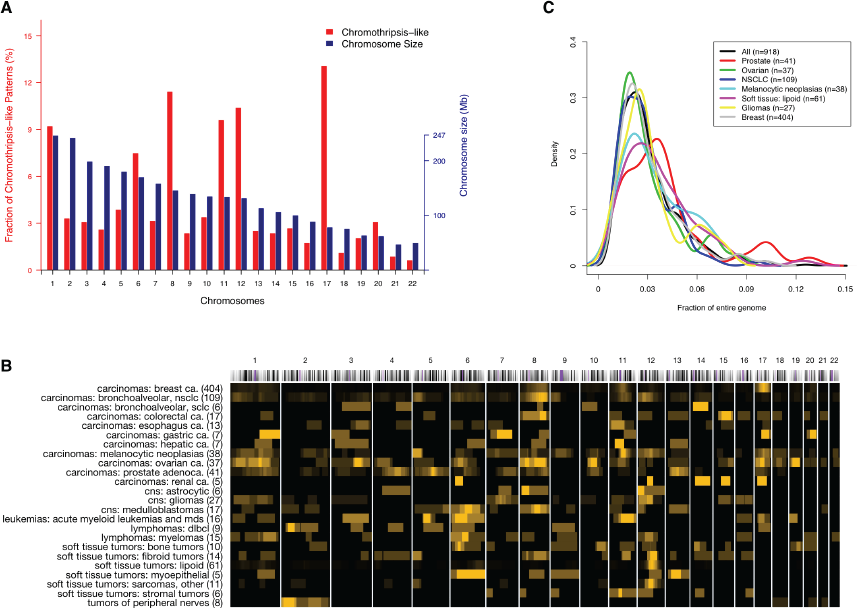
Frequency and CNA coverage length distribution of CTLP regions in the genome. (A) Red and blue bars indicate CTLP frequency in percent of all CTLP and chromosome size in megabases, respectively. (B) Local distribution of CTLP regions among diagnostic groups. Each row represents a cancer type and each column represents a chromosome. We use a black-to-yellow gradient for representing CTLP frequencies ranging from lowest to highest, normalized for each row. The numbers in brackets indicate the number of cases. Groups with at least 5 CTLP cases are shown. (C) Distribution of CTLP events as fractions of the affected genomes, represented as density plots for common cancer types. The fractions for each sample have been calculated as sum of genome bases chromosomes 1-22, divided by the genomic length of CTLP regions as identified through our scanning approach.

In our study, genomic projection of regional CTLP frequencies revealed their heterogeneous distribution in different cancer types (Figure 2B). The total length of fragmented genomic regions (CNA level and interspersed normal segments) accounted for 1%-14% of the corresponding genomes (Figure 2C). The large size of our input data set, resulting in high number of CTLP cases, permitted an investigation of the frequency and genomic distribution of these patterns in different cancer types. Our input samples represented 65 “diagnostic groups”, as defined by a combination of ICD-O morphology and topography codes. The majority of samples (18,238) came from 50 diagnostic groups, each represented by more than 25 arrays. We observed in total of 918 CTLP events across all 18,394 cases, representing an overall ∼5% prevalence. The 17 diagnostic groups represented by at least 45 cases, and having frequencies higher than 4% (CTLP high) are listed in Table 2 (full list in Additional file 5: Table S5).

**Table 2.** Frequency of chromothripsis-like patterns among cancer types.

| Cancer type | Chromothripsis-like cases |  |  |  | Input cases | Frequency (95% confidence interval) |
| --- | --- | --- | --- | --- | --- | --- |
|  | Oligo > 200K | Oligo ≤ 200K | BAC or cDNA | Total |  |  |
| Soft tissue tumors: lipoid | 49 | 12 | 0 | 61 | 114 | 53.5% (44%-62.8%) |
| Soft tissue tumors: fibroid tumors | 14 | 0 | 0 | 14 | 59 | 23.7% (14%-36.9%) |
| Soft tissue tumors: sarcomas, other | 9 | 0 | 2 | 11 | 48 | 22.9% (12.5%-37.7%) |
| Carcinomas: breast ca. | 247 | 99 | 58 | 404 | 3652 | 11.1% (10.1%-12.1%) |
| Carcinomas: esophagus ca. | 13 | 0 | 0 | 13 | 135 | 9.6% (5.4%-16.2%) |
| Carcinomas: bronchoalveolar, NSCLC | 78 | 29 | 2 | 109 | 1164 | 9.4% (7.8%-11.2%) |
| Soft tissue tumors: bone tumors | 7 | 3 | 0 | 10 | 123 | 8.1% (4.2%-14.8%) |
| Carcinomas: bronchoalveolar, SCLC | 3 | 3 | 0 | 6 | 90 | 6.7% (2.7%-14.5%) |
| Carcinomas: prostate adenoca. | 1 | 40 | 0 | 41 | 653 | 6.3% (4.6%-8.5%) |
| CNS: CNS PNET | 4 | 0 | 0 | 4 | 65 | 6.2% (2%-15.8%) |
| Carcinomas: melanocytic neoplasias | 31 | 1 | 6 | 38 | 621 | 6.1% (4.4%-8.4%) |
| Soft tissue tumors: myoepithelial | 3 | 0 | 2 | 5 | 85 | 5.9% (2.2%-13.8%) |
| Carcinomas: ovarian ca. | 31 | 5 | 1 | 37 | 801 | 4.6% (3.3%-6.4%) |
| Carcinomas: gastric ca. | 1 | 5 | 1 | 7 | 160 | 4.4% (1.9%-9.2%) |
| CNS: gliomas | 14 | 13 | 0 | 27 | 669 | 4% (2.7%-5.9%) |
| Soft tissue tumors: stromal tumors | 5 | 1 | 0 | 6 | 151 | 4% (1.6%-8.8%) |
| CNS: medulloblastomas | 13 | 4 | 0 | 17 | 430 | 4% (2.4%-6.4%) |

Only cancer types with input cases ≥ 45 and frequency ≥ 4% are shown

The initial study by Stephens *et al.* hypothesized that chromothripsis has a high incidence in bone tumors [9]. Notably, several soft tissue tumor types appeared in our “CTLP high” frequency set (6 out of 17), including the 3 types with the highest scores. Moreover, the high prevalence of CTLP in soft tissue tumors was reflected in the ICD-O specific frequencies (Additional file 6: Table S6). The genesis and/or effect of multiple localized chromosomal breakage-fusion events may be related to specific molecular mechanisms in those tumor types. Notably, gene fusions are well-documented recurring events in sarcomas [42], in contrast to most other solid tumors, and a local clustering of genomic re-arrangements had been previously reported for liposarcomas [43]. So far, more than 40 fusion genes have been recognized in sarcomas and treated as potential diagnostic and prognostic markers [42]. Possibly, the double-strand breaks and random fragment stitching events in chromothripsis-like events promote the generation of oncogenic fusion genes [9]. Further sequencing-based efforts will be needed to identify the true extent of fusion gene generation and to elucidate their functional impact in chromothripsis-like cases.

### Genomic context of chromothripsis-like events

It has been hypothesized that chromothripsis is a one-off cellular crisis generating a malignant clone in a very short time [9,44]. However, in many of the CTLP samples in our study, highly fragmented chromosomal regions were embedded in larger CNA regions showing variations in patterns and overall extent (Figure 3A). To test whether CTLP generating events are associated with overall genomic instability, we examined the extent of all copy number imbalances detected in our dataset. Comparing the 918 CTLP positive arrays with the remainder of 17,476 CTLP negative arrays, we found that CTLP samples tended to have higher proportions of CNA coverage in their genomes (*p* < 2.2×10^−16^; Kolmogorov-Smirnov test) (Figure 3B,C). This indicated that chromothripsis-like events frequently co-occur with other types of copy number aberrations. Plausible and non-exclusive explanations could be that CTLP might frequently arise due to previously established errors in the maintenance of genomic stability, or that chromothriptic aberrations involving genomic maintenance genes may predispose to the acquisition of additional CNA. For those frequent cases exhibiting additional non-CTLP CNA events, their possible contribution to oncogenesis has to be considered when modeling the role of chromothripsis-like events in cancer development.

**Figure 3.**
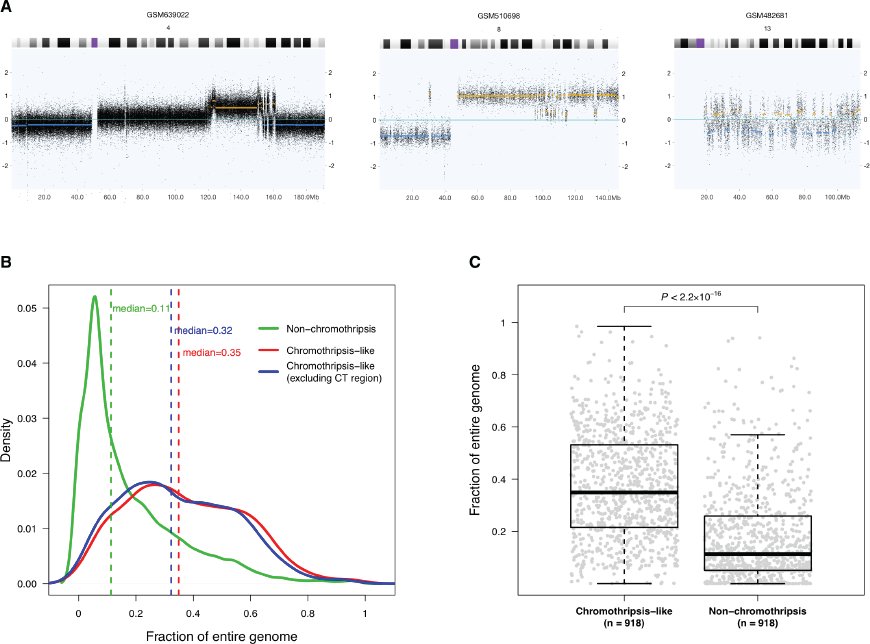
Genomic context of CTLP events. (A) Example copy number profiles of chromosomes with changes suggestive of chromothripsis. In these examples, chromosomal fragmentation events are related to other types of copy number aberrations, and exhibit different combination patterns. The x-axis indicates genomic locations in Mb, and the y-axis is the log2 value of probe signal intensity. Yellow and blue lines represent called genomic gains and losses respectively. (B) Distribution of CNAs as fraction of the genome, compared between CTLP and non-CTLP cases. CT, chromothripsis-like. (C) Distribution of CNA fractions for individual samples. For the non-CTLP group, 918 samples were randomly chosen from the total set of 17,476 cases, to generate an equally sized comparison. The *p*-value, indicating significant difference between the genome fraction distributions of two groups, is based on a Kolmogorov-Smirnov test. The fractions for each sample have been calculated as sum of genome bases chromosomes 1-22 divided by the sum of all CNAs in the sample (with and without CTLP regions).

### Potential mechanisms for chromosome shattering

While the mechanism(s) responsible for the generation of chromothripsis remain elusive, a number of studies have proposed hypotheses including ionizing radiation [9], DNA replication stress [45], breakage-fusion-bridge cycles [9,23,46], premature chromosome compaction [47], failed apoptosis [48,49] and micronuclei formation [50]. Some of these proposed mechanisms are associated with features which could be addressed in our study.

In our dataset, although most (76%) CTLP cases presented single chromosome CTLP events, in approximately 24% CTLP affected at least 2 chromosomes (Figure 4A). For certain candidate mechanisms, e.g. micro-nucleus formation due to mitotic delay, this observation would imply more than one event, whereas the observation appears compatible with e.g. an aborted apoptosis process.

**Figure 4.**
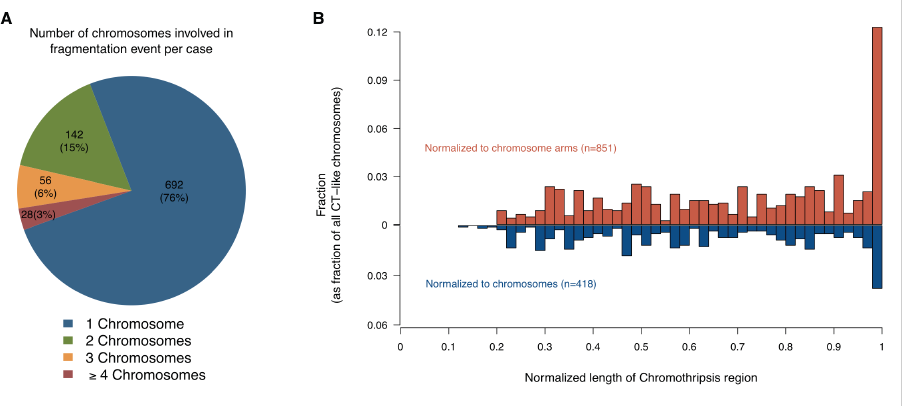
The distribution of CTLP regions in terms of chromosome number and length. (A) The number of chromosomes affected by CTLP per sample. The numbers outside and inside the brackets are number and percentage of CTLP samples respectively. (B) Length distribution of CTLP regions normalized to chromosome or chromosome arm lengths. For each chromosome, regions restricted on a single arm were normalized to arm lengths (red bars), otherwise were normalized to chromosome lengths (blue bars). More than 10% of all CTLP events involve whole chromosome arms.

For relating to cytogenetic aberration mechanisms, an additional parameter explored by us was the extent of CTLP regions when normalized to their respective chromosomes. Affected regions were classified into the categories “arm-level” (≥ 90% arm length), “chromosome-level” (≥ 80% chromosome length) or “localized” (Figure 4B). Arm-level CTLP events were observed with a relatively high frequency (∼19%). In the arm-level patterns, the CTLP rearrangements were concentrated in one chromosome arm, with the other arm of the same chromosome remaining normal or showing isolated CNA. Since arm-level events involve both peri-centromeric and telomeric regions, cytogenetic events involving these chromosomal structures present themselves as possible causative mechanisms.

Notably, one model that closely conforms to this pattern involves breakage-fusion-bridge cycles [9,23,46,47,51–54]. In general, such cycles start with telomere loss and end-to-end chromosome fusions. When the dicentric chromosomes are formed and pulled to opposite poles during anaphase, a double-strand DNA break acts as starting point for the next cycle. Chromosomal rearrangements would gradually accumulate during the additional cycles, and should be concentrated in one chromosome arm, particularly near the affected telomere. In our dataset, up to 44% of all CTLP chromosomes involved telomere regions. We performed simulations to explore whether this telomere enrichment could be explained by chance. In brief, for each sample, we retained the location of CTLP region in the genome and shuffled the telomere position of each chromosome while keeping the length of each chromosome constant. In contrast to the actual observations, the simulation did not result in telomeric CTLP enrichment (*p* < 0.0001; 10,000 simulations; see Methods). CTLP generation through breakage-fusion-bridge cycles would be a viable candidate hypothesis compatible both with the statistically significant telomere enrichment and the high proportion of arm-level pulverization. However, for arm-level CTLP events centromere-related instability mechanisms should also be considered for future discussions.

### Clinical implications

Based on clinical associations of “chromothripsis” patterns, it has been claimed that these events may correlate with a poor outcome in the context of the respective tumor type [14,25,55]. In our meta-analysis, we explored a general relation of CTLP with clinical parameters, across the wide range of cancer entities reflected in our input data set. Clinical data was collected from GEO and from the publications of the respective series (Additional file 2: Table S2 and Additional file 1: Table S7) and parameters available for at least 1,000 cases were considered. From our dataset, CTLP seemed to occur at a more advanced patient age as compared to non-CTLP samples (Figure 5A) [23]. CTLP mainly occurred at stage II and III (70%), which was significantly different from the stage distribution of total samples (55.2%) (*p* = 0.0149; Chi-square test) (Figure 5B). No difference of grade distribution was observed in our dataset (*p* = 0.425; Chi-square test) where CTLP samples showed a predominance for grades 2 and 3, similar to the bulk of all samples (∼80%). We also found that CTLP was overrepresented in cell lines compared to primary tumors (*p* < 2.2×10^−16^; two-tailed Fisher’s exact test).

**Figure 5.**
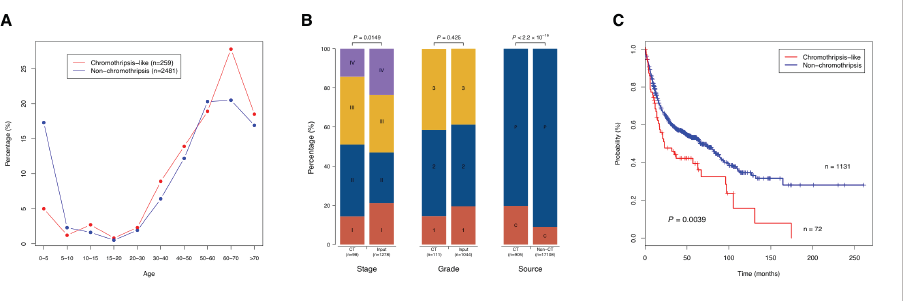
Clinical perspective on CTLP events. (A) Distribution of CTLP percentage versus patient age. (B) Distribution of sample stage, grade and source between CTLP and input dataset or non-CTLP cases. *p*-values are derived from Chi-square test (stage and grade) or two tailed Fisher’s exact test (source). P, primary tumor; C, cell line. (C) Kaplan-Meier survival curves for CTLP versus non-CTLP cases. The *p*-value is based on log-rank test.

For a subset of 1,203 patients, we were able to determine basic follow-up parameters (follow-up time and survival status). For 72 of these individuals, CTLP was detected in their tumor genomes. Notably, patients with CTLP survived a significantly shorter time than those without this phenomenon (*p* = 0.0039; log-rank test; Figure 5C). Note that this analysis was based on a sample of convenience averaged over cancers, stages and grades. If we break down this dataset by cancer type, the numbers are not large enough to provide statistical confidence (Additional file 1: Figure S4). While the cancer type independent association of CTLP patterns and poor outcome is intriguing, potential clinical effects of chromothripsis-like genome disruption should be evaluated in larger and clinically more homogeneous data sets.

### Sensitivity of array platforms for detection of chromothripsis-like patterns

Presumed chromothripsis events have been reported from genomic datasets generated through different array and sequencing based techniques (see table 1). We performed an analysis of the platform distribution of our CTLP samples, to estimate the detection bias among various genomic array platforms. As the resolution of a platform depends both on type and density of the probes on an array, we divided the platforms into 4 groups according to their probe numbers and techniques (BAC/P1, DNA/cDNA, oligonucleotide ≤ 200 K and oligonucleotide > 200 K). Although CTLP were detected by all types of genomic arrays, a higher fraction of CTLP samples was found using data from high resolution oligonucleotide arrays (Figure 6), possibly due to increased sensitivity related to higher probe density. Indeed, when performing platform simulations, the sensitivity of CTLP detection improved with increasing probe numbers (Additional file 1: Figures S5 and S6; see Methods). According to these simulations, array platforms consisting of more than 250k probes should be preferred when screening for CTLP events. Since our analysis relied on a variety of array platforms, we can assume that the overall prevalence of CTLP in cancer is higher than our reported 5% of samples.

**Figure 6.**
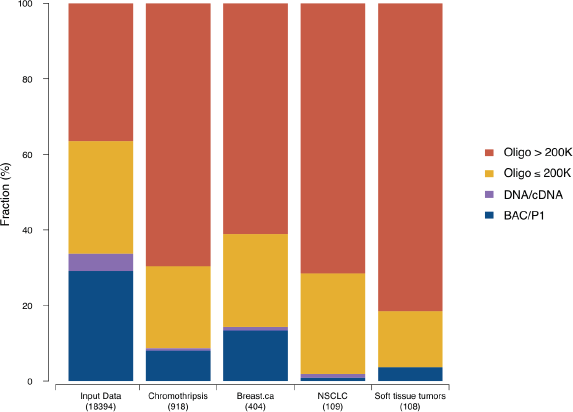
Platform distribution based on different resolutions and technology types. Different analysis groups are shown, including the whole input dataset, inferred CTLP cases and three cancer types. The larger fraction of high-density oligonucleotide arrays in samples with CTLP compared to the overall platform distribution indicates an increased sensitivity of these platforms for CTLP events. Oligo, oligonucleotide; NSCLC, Non-small cell lung cancer.

## Discussion

The description of the “chromothripsis” phenomenon has initiated a vital discussion about clustered genomic aberration events and their role in cancer development [52,55,56]. While chromothripsis *senso stricto* has been characterized as a type of focally clustered genomic aberrations generated in a one time cellular event and being limited to a defined set of copy number states [15], other operational definitions have been employed based on clustered aberrations [7,16,23,45,55,57]. It seems likely that some of the previous discussions of “chromothripsis” referred to a number of underlying event types, all resulting in localized genome fragmentation and re-assembly events. For instance, DNA double strand break and end-joining-mediated repair may result in a restricted number of copy number levels, whereas aberrant replication based mechanisms will lead to a more diverse set of copy number aberrations [45,55]. Here, we introduce the term “chromothripsis-like patterns” (CTLP) when referring to clustered genomic events, to accommodate both common labelling and presumed biological variability of clustered genomic copy number aberrations.

At this time, due to the lack of sufficiently large number of cancer data sets from whole-genome sequencing analyses, a meta-analysis of “strict” chromothripsis cases is not feasible. We have followed a pragmatic approach to quantify the occurrence of CTLP from genomic array data sets. In our algorithm, we implemented the two most significant features shared by different operational chromothripsis definitions, namely copy number status changes and breakpoints clustering, which can be well measured by array based technologies. Previous studies provided various algorithms to detect “chromothripsis” events [9,15,58]. However, besides its application to an extensive data set, the specific advantage of our method presented here is its ability to detect regions of shattering with limited influence from the varying sizes of affected chromosomes. Since the step length of our scanning window is 5 Mb, theoretically the detected CTLP regions are within an accuracy of ± 5 Mb. Note that the performance of this algorithm may be influenced by poor quality arrays, especially those with highly scattered and unevenly distributed probe signal intensities.

In this study, we identified 918 CTLP-containing genome profiles, based on an analysis of copy number aberration patterns from 22,347 oncogenomic arrays and representing 132 cancer types. Despite the inherent limitations of such a meta-analysis approach, we were able to provide several new insights regarding the distribution of clustered genomic copy number aberrations and to produce a comprehensive estimate of CTLP incidence in a large range of cancer entities.

In our analysis, CTLP exhibited an uneven distribution along tumor genomes, with disease related local enrichment. These “CTLP dense” chromosomal regions may reveal associations between disease related cancer associated genes and molecular mechanisms behind genome shattering events. This potential correlation is exemplified by the prevalence of mutant *TP53* in “chromothriptic” Li-Fraumeni syndrome associated Sonic-Hedgehog medulloblastomas [23]. As the extent of CTLP related deletions of the *TP53* locus indicates, CTLP related gene dosage changes may predispose to double-hit effects on specific tumor suppressors. In contrast, we found regional enrichment for CTLP with pre-dominant copy number gains on chromosomes 8, 11 and 12. In the initial study, chromosome 8 shattering was found in a small cell lung cancer cell line [9]. This event contained the *MYC* oncogene, which had be shown to be amplified in 10-20% of small cell lung cancers [59]. Moreover, strong overexpression of *MYC* involved in a “chromothripsis” region was also detected in a neuroblastoma sample [25]. In a study of colorectal tumors, chromosomes 8 and 11 were involved in concurrent pulverization events with generation of fusion genes, involving e.g. *SAPS3* and *ZFP91* [18]. In a study on hepatocellular carcinoma, *CCND1* amplification was embedded within a “chromothriptic” event on chromosome 11 [24]. Therefore, the overall uneven distribution of CTLP may point to specific driver mutations that contribute to CTLP generation, and/or to a class of cancer promoting mutations based on regional genome shattering events.

When comparing cancer types, we observed a high CTLP prevalence in a limited set of entities, particularly in among soft tissue tumors. This finding supports and improves upon a previous prediction of particularly high “chromothripsis” rate in bone tumors [9]. Also, the uneven distribution of CTLP is a strong indicator for a disease related selection of specific genomic aberrations, supporting their involvement in the oncogenetic process.

In the initial study, the authors stated that chromothripsis could be a one-off cataclysmic event that generates multiple concurrent mutations and rearrangements [9]. However, the role of chromothripsis in terms of “shortcut” to cancer genome generation is still elusive. We note that additional and complex non-CTLP genome re-arrangements exist in the majority of CTLP samples. The number and uneven distribution of affected chromosomes in CTLP supports the biological heterogeneity of cancer samples with CTLP containing genome profiles. Furthermore, the normalized spatial distribution of shattered chromosomal regions, as well as the observed significant overlap between telomere and pulverized regions is supportive of breakage-fusion-bridge cycles as one of the mechanisms acting in a subset of samples. Further efforts are needed to investigate the temporal order of chromothripsis and non-chromothripsis events in complex samples, and to substantiate the existence of a dichotomy between “one-off” chromothripsis and other classes of localized genome shattering events, all resulting in clustered genomic copy number aberrations.

In our associated clinical data, CTLP were related to more advanced tumor stages and overall worse prognosis when compared to non-CTLP cases. One possible explanation is that the numerous concurrent genetic alterations induced by genome shattering events disturb a large number of genes and contribute to more aggressive tumor phenotypes. By themselves, these observations do not differentiate whether CTLP arise as a early events promoting aggressive tumor behavior with fast growth rates and reduced response rates to therapeutic interventions; or whether this observation relates to underlying primary mutations predisposing to genomic instability, aggressive clinical behavior and CTLP as a resulting epiphenomenon. Interestingly, the high rate of *TP53* involvement by itself would support both possibilities for this gene, i.e. chromothripsis as result of *TP53* mutation as well as chromothriptic events with *TP53* locus involvement promoting an aggressive clinical behavior.

From Table 1 we may notice that the array based technologies are, in general, less sensitive than whole-genome sequencing data for calling chromothripsis-like events. This is partly due to the very limited ability of most array platforms to detect balanced genomic aberrations, such as inversions and translocation events. In the future, the accumulation of large-scale sequencing data should be able to provide further insights into localise genome shattering events.

## Conclusions

CTLP represent a striking feature occurring in a limited set of cancer genomes, and can be detected from array based copy number screening experiments, using biostatistical methods. The observed clustered genomic copy number aberrations may reflect heterogenous biological phenomena beyond a single class of “chromothripsis” events, and probably vary in their specific impact on oncogenesis. Fragmentation hotspots derived from our large-scale data set could promote the detection of markers associated with genome shattering, or may be used for assigning disease related effects to CTLP-induced genomic events.

## Methods

### Genome-wide microarrays and data preparation

In this study, we screened 402 GEO series [36], encompassing 22,347 high quality genomic arrays (Additional file 2: Table S2). All selected arrays were human cancer samples hybridized onto genome-wide array platforms. The normalized probe intensities, segmented data and quality information were obtained from the arrayMap database, which is a publicly available reference database for copy number profiling data [37]. In brief, the annotated data was obtained by the following processing pipeline: for Affymetrix arrays, the aroma.affymetrix R package was employed to generate log2 scale probe level data [60]; for non-Affymetrix arrays, available probe intensity files were processed; CBS (Circular Binary Segmentation) algorithm [61] was performed to obtain segmented copy number data. The probe locations were mapped on the human reference genome (UCSC build hg18). In the case of technical repeats (e.g. one sample was hybridized on multiple platforms), only one of the arrays was considered for analysis (preferably with the highest resolution and/or best overall quality). The array profiling can be visualized through the arrayMap website.

### Scan-statistic based chromothripsis-like pattern detection algorithm

To detect chromothripsis-like cases, we formulated an algorithm identifying clustering of copy number status changes in the genome. Several parameters were considered to define the alteration of copy number status:

i. The thresholds of log2 ratio for calling genomic gains and losses. These values were array specific and stored in arrayMap database. For each array, the thresholds were obtained from related publications or empirically assigned based on the log2 ratio distribution.
ii. The intensity distance between adjacent segments. Due to local correlation effects between probes or the existence of background noise, the segmentation profiles occasionally exhibit subtle striation patterns. This pattern is constituted with a large number of small segments, which is unlikely to be a biological phenomenon. To reduce artificial copy number status change, the distance of signal intensity between adjacent segments was used as a threshold, and defined here as the sum of the absolute values to call gains and losses. If the distance of two adjacent segments differed by less than this threshold, the copy number status change was not considered.
iii. Segment size. The resolution of a platform depends on the density of probes on the array. One of the platforms with the highest density in our dataset is Affymetrix SNP6, which contains 1.8 million polymorphic and non-polymorphic markers with the mean inter-marker distance of 1.7 kb. It provides a practical resolution of 10 to 20 kb. Therefore, in this study, segments smaller than 10 kb were removed.

In order to identify clustering of copy number status changes, a scan-statistic likelihood ratio based on the Poisson model was employed [39]. In our implementation, a fixed-size window was moved along the genome and for each window the likelihood ratio was computed from observed and expected copy number status change times. Let *G* be the genome represented linearly, and *W* is a window with fixed size. As the window *W* moves over *G*, it defines a collection of zones *Z*, where *Z* ⊂ *G*. Let *n_W_* denotes the observed copy number status change times in window *W*, and *n_G_* the total number of observed status change in *G*. *μ*_*W*_ is the expected status change times in window *W*, and is calculated as *W/G*×*n*. The likelihood function is expressed as

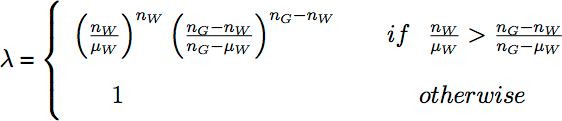

This function detects the zone that is most likely to be a cluster.

Due to lack of prior knowledge about the size of *W*, we predefined a series of window sizes from 30 Mb to 247 Mb (Additional file 1: Table S8), which were based on chromosome sizes. The scanning process was repeated for the series of window sizes for each sample. When *W* moved over *G*, the step length was set to 5 Mb, and there was no overlap between different chromosomes in window *W*. In this way, for each genome, the collection of *Z* contained 4,414 windows in various sizes. The window that maximized the likelihood ratio defined the most probable CTLP region. Thus it can detect both the location and the size of the cluster. When analyzing the complete input dataset, the window with the highest likelihood ratio was selected as a candidate of chromothripsis for each chromosome of the 22,347 arrays. The R script for detecting CTLP cases can be provided upon request.

### Analysis of fragment enrichment in telomere region

Telomere positions were simulated to test the DNA fragment enrichment. For each case, the CTLP region was kept at its location in the genome. Locations of chromosome terminals were randomly selected while the length of each chromosome was kept. A genomic interval of 5 Mb from the chromosome terminal was considered as the telomere region. The simulation was performed 10,000 times.

### Simulation of platform resolution

The 15 Affymetrix SNP6 CTLP chromosomes in the training set were used for simulation. For each genome, a certain number of probes were randomly chosen from the original probe set. These probes generally represented the profile that the same sample was hybridized on a platform with corresponding resolution. Then the CTLP pattern detection algorithm was applied on the simulated arrays, and the number of cases that passed the thresholds were recorded.

### Statistical testing

The significance in the number of CTLP cases with *TP53* loss in comparison to those in non-CTLP cases was assessed using two-tailed Fisher’s exact test. We performed a Kolmogorov-Smirnov test to compare the distributions of copy number aberration proportions in the genome between CTLP and the other cases. The Chi-square test was used to assess the significance in the distribution of both patient stage and grade in CTLP and the whole input dataset. The associations between the number of cell lines in CTLP and non-CTLP cases were tested by two-tailed Fisher’s exact test. The difference in the survival curves between two subgroups was evaluated by the log-rank test.

## Authors’ Contributions

HC, MDR and MB conceived and designed the experiments. HC and NK analyzed the data. HC, NK, HCB, CvM, MDR and MB contributed reagents/materials/analysis tools. All authors contributed to draft the manuscript and approved the final manuscript.

## Acknowledgements

The authors would like to thank Henrik Bengtsson and Ni Ai for useful discussions.

## Additional Files

**Additional file 1: Supplementary figures and tables.**

**Figure S1:** Scatter plot of the training set.

**Figure S2:** The positive training set and CTLP detection algorithm performances.

**Figure S3:** Scatter plot of CTLP candidates.

**Figure S4:** Kaplan-Meier survival curves for CTLP versus non-CTLP cases in specific cancer types.

**Figure S5:** An example of the platform resolution based simulation using data from an Affymetrix SNP6 array (∼1.8 million probes).

**Figure S6:** CTLP detection sensitivity of simulated platform resolutions.

**Table S1:** Overview of input dataset.

**Table S7:** Demographic and clinico-pathologic characteristics of input and CTLP samples.

**Table S8:** Sizes of sliding windows for the scan-statistic based algorithm.

### Additional file 2

**Table S2.** Input dataset of high quality arrays with probe level raw data.

### Additional file 3

**Table S3.** Training set collected from published chromothripsis cases.

### Additional file 4

**Table S4.** Detected chromothripsis-like cases.

### Additional file 5

**Table S5.** Frequency of chromothripsis-like pattern among diagnostic groups.

### Additional file 6

**Table S6.** Frequency of chromothripsis-like pattern among ICD-O codes.

